# Evolutionary Changes in Left-Right Visceral Asymmetry in *Astyanax* Cavefish

**DOI:** 10.1101/2020.05.15.098483

**Authors:** Li Ma, Mandy Ng, Janet Shi, Aniket V. Gore, Daniel Castranova, Brant M. Weinstein, William R. Jeffery

## Abstract

Vertebrates show conserved left-right (L-R) asymmetry of internal organs controlled by Nodal-Pitx2/Lefty signaling [1-3]. Modifications in L-R asymmetry occur in mutants [4] and rarely in humans [5], but little is known about natural L-R changes during evolution. Here we describe changes in L-R asymmetry in *Astyanax mexicanus*, a teleost with ancestral surface (surface fish) and derived cave (cavefish) morphs [6]. In teleosts, Nodal-Pitx2 signaling is activated in the left lateral plate mesoderm (LPM), the cardiac tube jogs to the left and loops to the right (D-looping), and the liver and pancreas form on opposite sides of the midline. Surface fish show conventional L-R patterning, but cavefish can show Nodal-Pitx2 expression in the right LPM or bilaterally, left (L)-looping hearts, and reversed liver and pancreas asymmetry, and these reversals have no effect on survival. The Lefty1 Nodal antagonist is expressed along the surface fish and cavefish midlines, but expression of the Lefty2 antagonist is absent in the LPM of most cavefish embryos, suggesting a role for *lefty2* (*lft2*) in changing organ asymmetry. Although CRISPR-Cas9 *lft2* editing affected D-looping in surface fish, the cavefish *lft2* gene showed no coding mutations, and was expressed normally during cavefish gastrulation, suggesting downregulation by regulatory changes. Reciprocal hybridization, the fertilization of cavefish eggs with surface fish sperm and *vice versa*, indicated that the change in cavefish L-R asymmetry is a maternal genetic effect. Our studies reveal natural changes in internal organ asymmetry during evolution and introduce *A. mexicanus* as a new model to study the underlying mechanisms.

## Results and Discussion

Bilaterally symmetric animals generally show superficial symmetry of external structures but distinct asymmetries in many internal organs. During vertebrate development visceral organs such as the heart and gut tubes bend asymmetrically and develop at predetermined positions favoring the left or right side of the midline [1-4]. This direction of left-right asymmetry is conserved in amphibians, teleosts, birds and mammals [5, 6]; however little is known about the possibility of internal organ asymmetry changes during vertebrate evolution. The teleost *A. mexicanus*, which consists of an ancestral surface-dwelling morph (surface fish) and many derived cave-dwelling morphs (cavefish), is a model system for studying the evolution of development [7]. In cavefish adults, asymmetry transcends the internal organs and is also present in craniofacial structures, such as the jaws, skull, and cranial neuromasts [8, 9]. However, evolutionary changes in L-R patterning of cavefish internal organs have not been investigated during embryogenesis.

### Changes in Internal Organ Asymmetry in Cavefish

To understand whether internal organ asymmetry can be changed in cavefish, we investigated asymmetric development of the heart, liver, and pancreas. The heart primordium is the first organ precursor to show L-R asymmetry in vertebrate embryos [10]. The cardiac tube forms in LPM along the ventral midline, then jogs to the left and loops to the right, giving rise to an S-shaped organ. The direction of cardiac jogging and looping was compared in surface fish, Pachón cavefish (PA-CF), and Tinaja cavefish (TI-CF). At 1.5 days post-fertilization (dpf), more than 98% of surface fish larvae showed left jogging cardiac tubes. In contrast, only 60-75% of cardiac tubes jogged to the left in PA-CF and TI-CF larvae, and the remainder showed either right jogging or no jogging (Fig. S1). To determine the direction of heart looping, surface fish and cavefish embryos were stained with a myosin heavy chain antibody [11,12] (Fig. 1A, B). At 3 dpf, more than 95% of surface fish larvae showed D-looping hearts, whereas only 75% of PA-CF larvae and 86% of TI-CF larvae had D-looping hearts, and the rest showed either L-looping hearts (13% and 4% respectively; Supplementary Movie) or non-looping hearts (7% or 9% respectively). The proportion of modified heart looping varied among different PA-CF families: some families (e.g. F61) exhibited almost 40% abnormal heart looping, others exhibited only moderate levels of altered heart looping (e.g. F72), and still others (e.g. F64) showed mostly D-looped hearts (Fig. 1C). The reason for this variation is unknown but could be a result of incomplete fixation of the allele(s) controlling L-R determination. The results indicate that conventional heart asymmetry can be changed in cavefish, most prominently in certain PA-CF families.

**Figure 1.**
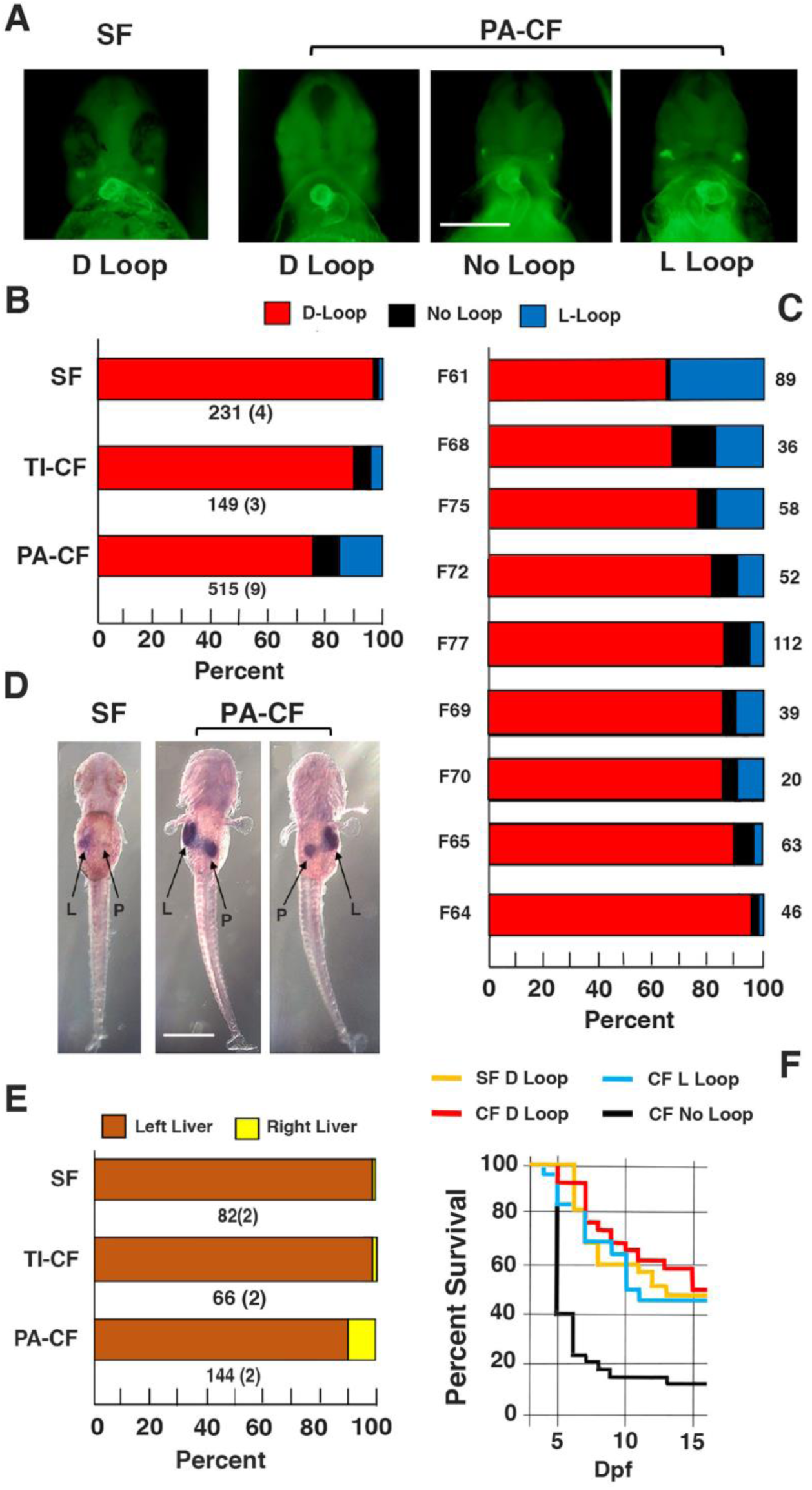
Changes in internal organ asymmetry in cavefish. A, B. The direction of heart looping is changed in cavefish. A. Surface fish (SF) with D-looping heart and Pachón cavefish (PA-CF) with D-looping, no looping, or L-looping hearts. Embryos stained with MF-20 antibody viewed from the ventral side at 3 dpf. Scale bar: 250µM; magnification is the same in all frames. B. Bar graphs showing the percentage of heart looping types in surface fish, PA-CF, and TI-CF. Bottom of bars: Total number of fish assayed (number of different families). C. Bar graphs showing the differences in the proportion of heart looping changes in PA-CF families. Family (F) numbers are shown on the left of the bars. Number of assayed fish are indicated at the right of each bar. D, E. Liver and pancreas laterality in surface fish and cavefish determined by *cbsa* expression. D. *In situ* hybridization showing *cbsa* expression in liver positioned on the left and pancreas positioned on the right of the midline in surface fish, and normal and reversed liver-pancreas laterality in PA-CF at 3.5 dpf. All views from the dorsal side. L: liver. P: pancreas. Scale bar in is 500µm; magnification is the same in all frames. E. Bar graphs showing the percentage of livers positioned on the left or right side in SF, TI-CF, and PA-CF). Total number of *in situ* hybridized larvae is shown at the bottom of each bar, followed by the number of families analyzed in parentheses. F. Survival of cavefish with differences in heart laterality. Graph shows the percentage of surviving surface fish (SF) of an initial 100 larvae with D-looped hearts and PA-CF of an initial 75 larvae with D-looped hearts, an initial 38 larvae with non-looped hearts, and an initial 52 larvae with L-looped hearts on each day beginning at 3 days post-fertilization (dpf). Red, blue, and yellow survival profiles show no significant differences from each other. Black survival profile shows a significant difference (p < 0.000) compared to all other groups.

We next asked whether changes in L-R asymmetry are restricted to the heart or also occur in other cavefish internal organs. The teleost liver and pancreas form as tubular protrusions of the gut, which subsequently shift from the midline and continue to develop on the left or right sides of the body respectively [5]. The positions of liver and pancreas development were compared in surface fish, PA-CF, and TI-CF by *in situ* hybridization using the *cbsa* gene as a marker [13]. Consistent with the conventional L-R laterality in zebrafish [14], at 3 dpf the liver developed on the left and the pancreas on the right in more than 98% of surface fish larvae (Fig. 1D, E), and TI-CF larvae showed the similar liver-pancreas asymmetry (Fig. 1E). However, only 89% of PA-CF larvae showed the normal L-R relationship (Fig. 1D, E), and the liver developed on the right and the pancreas on the left in the remaining 11% of PA-CF larvae (Fig. D, E), showing that liver and pancreas development can also be reversed in PA-CF larvae. Thus, cavefish have evolved changes in both heart and endodermal organ asymmetry.

To determine whether changes in L-R asymmetry affect cavefish viability, we compared survival between surface fish and F61 PA-CF, the family showing the highest levels abnormally looped hearts (Fig. 1C). Cavefish larvae were separated into groups with D-looped, non-looped, and L-looped hearts at 3 dpf, and the number of living larvae in each group, and in surface fish controls, was followed for the next 14 days (Fig. 1F). The surface fish controls, all with D-looped hearts, showed about 50% survival, which is typical under our culturing conditions [13], and there were no differences between survival of surface fish or cavefish with D-looped hearts and cavefish with L-looped hearts (Fig. 1F). In contrast, survival was significantly lower in most cavefish without heart looping, and most of these larvae perished. The results indicate that complete reversal of heart looping does not affect survival in cavefish, although in most cases the absence of cardiac looping is lethal.

### Changes in the Nodal-Pitx2 Signaling Cascade in Cavefish

Nodal signaling begins in bilateral domains surrounding Kupffer’s vesicle (KV), the teleost symmetry breaking organizer [15], then during segmentation continues in the left LPM, where it activates *pitx2* expression, and the Nodal-Pitx2 cascade is suppressed in the right LPM. To address the possibility that alterations in L-R asymmetry of Nodal/Pitx2 signaling are associated with cavefish organ laterality, expression of the teleost *nodal* paralog *southpaw* (*spaw)* [16] and *pitx2* genes was compared by *in situ* hybridization in surface fish and cavefish embryos. F61 PA-CF (Fig. 1C) were used in these and the other experiments described below. At the 8-10 somite stage, *spaw* was expressed bilaterally around the KV in both surface fish and cavefish embryos (Fig. 2A-D), suggesting that the Nodal signaling begins normally on both sides of the L-R axis. Later in development, at the 18-25 somite stage, surface fish embryos expressed *spaw* and *pitx2* in the left LPM, but not in the right LPM (Fig 2E, I, J, N), indicative of normal Nodal-Pitx2 cascade asymmetry. However, *spaw* and *pitx2* were expressed in the left LPM in only 75% of cavefish larvae, whereas these genes were expressed in the right LPM in 20%, and bilaterally in 5%, of the cavefish embryos (Fig. 2F-I, K-N). Therefore, the left-sided asymmetry of Nodal-Pitx2 signaling is modified in conjunction with L-R axial changes in cavefish.

**Figure 2.**
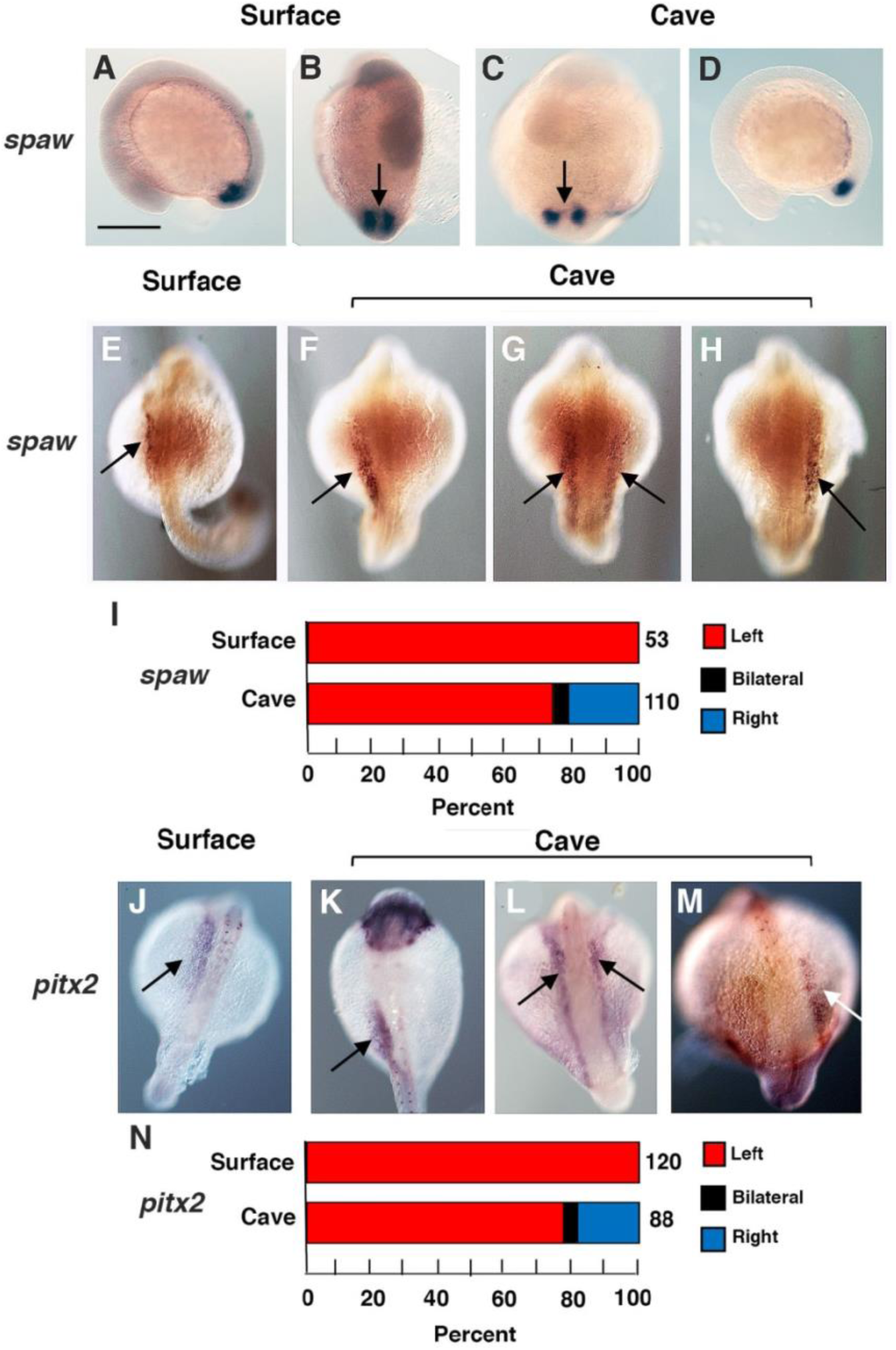
Changes in the Nodal-Pitx2 signaling cascade in cavefish. A-D., E-H. *In situ* hybridizations showing *spaw* expression around Kupffer’s vesicle (arrow) and in the lateral plate mesoderm (LPM) at the 10-13 somite (A-D) and 18-25 somite (E-H) stages in surface fish (A, B, E) and in cavefish (C, D; F-H) embryos. J-M. *In situ* hybridizations showing *pitx2* expression in the LPM of 25-30 somite surface fish (J) and cavefish (K-M) embryos. I, N. Bar graphs showing number of embryos with *spaw* (I) or *pitx2* (N) expression in the left, left and right, and right LPM in surface fish and cavefish embryos. Number of embryos analyzed is shown to the right of each bar. A, C. lateral views with anterior to the left. B, D, E-H, J-M. Dorsal views with anterior on the top. Unlabeled arrows: expression in LM. Scale bar in A: 100 µm; magnification is the same in all frames.

### Role of *Lefty* Genes in Cavefish Left-Right Asymmetry

To determine whether the Nodal autoregulatory system involving the *lefty* genes is functional in cavefish, we examined *lefty1* (*lft1*) and *lefty2* (*lft2*) expression in surface fish and PA-CF embryos. The Nodal-Pix2 cascade is restricted to the left LPM by the Lefty1 and Lefty2 barriers, which antagonize Nodal and prevent its ectopic expression in the right LPM [17-19]. *In situ* hybridization showed that *lft1* is expressed at similar levels along the midline of 18-23 somite surface fish and cavefish embryos (Fig. 3A-D), suggesting that the Lefty1 barrier remains intact in both morphs. In contrast, although *lft2* is expressed normally in the heart primordium of the left LPM in surface fish embryos, expression could not be detected in either the left or right LPM in the majority of cavefish embryos (Fig. 3F-H; Fig. S2). In about 5% of cavefish embryos, the *lft2* gene was expressed weakly and slightly earlier than in surface fish but only in the left LPM (Fig. 3G, H; S1). To substantiate these results, double staining *of lft1* and *lft2* was done, and all cavefish embryos stained positive for *lft1*, but none showed *lft2* staining (Fig. S3). To confirm *lft2* downregulation, the levels of *lft2, lft1*, and the *nodal related 2*(*ndr2*) mRNA were quantified by qPCR (Fig. 3E). A significant decrease in *lft2* levels was seen in cavefish relative to surface fish embryos, and *lft1* and *ndr2* levels were also increased in cavefish embryos, although they did not reach significance (Fig. 3E). We conclude that *lft2* expression is strongly downregulated in the cavefish LPM, suggesting that the Lefty1 barrier to Nodal diffusion may be disrupted.

**Figure 3.**
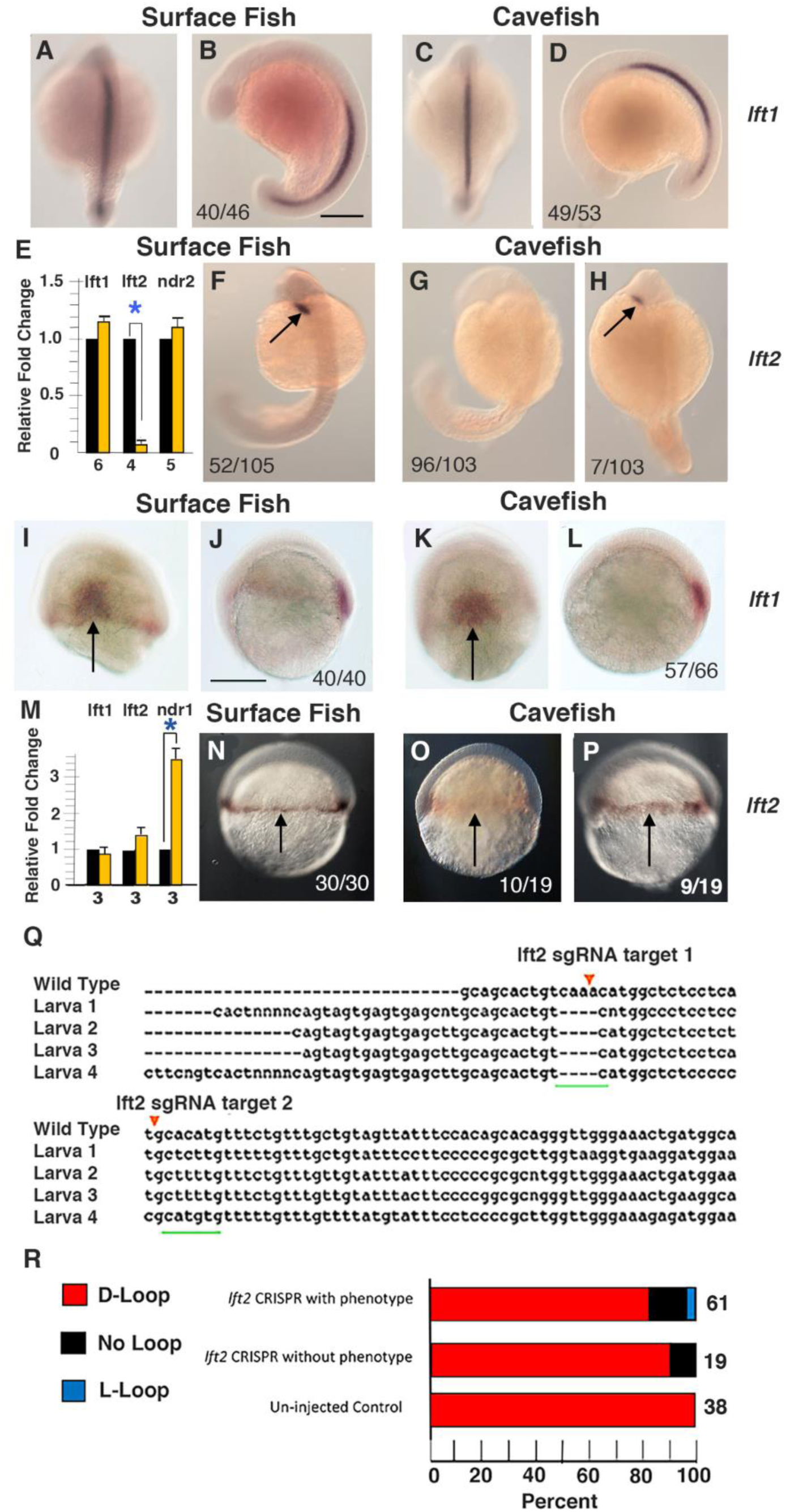
Role of *lefty* genes in cavefish left-right asymmetry. A-D. *In situ* hybridization showing *lft1* expression at the dorsal midline in 10-13 somite surface fish (A, B) and cavefish (C, D) embryos. A, C: Dorsal views with anterior on the top. B, D. Lateral views with anterior on the left. F-H. *In situ* hybridization showing *lft2* expression in the cardiac fields (arrows) of the lateral plate mesoderm (LPM) in 18-25 somite surface fish and cavefish embryos. Dorsal views. Most cavefish embryos did not show detectible *lft2* expression in the LPM (G), but in a small number of cavefish embryos *lft2* was expressed weakly compared to surface fish (H). Scale bar in B: 100µm; magnification is the same A-D and F-H. I-L. *In situ* hybridization showing *lft1* expression in the shield region (arrows) of surface fish (I, J) and cavefish (K, L) embryos at 50% epiboly. I, K. Dorsal views. J, L: Lateral views. N-P. *In situ* hybridization showing *lft2* expression in the germ ring (arrows) of surface fish (N) and cavefish (O, P) embryos at 50% epiboly. Numbers in frames indicate proportion of embryos with indicated expression pattern. Scale bar in J: 150 µm; magnification is the same in I-L and N-P. E, M. Bar graphs showing qPCR analysis of relative mRNA fold changes in cavefish compared to surface fish at the 25 somite (E) and 40-50% epiboly (M) stages. Black columns: surface fish. Yellow columns: cavefish. Error bars: SEM. Number of replicates shown at the bottom of the columns. Asterisks: significance at p < 0.01. Q-R. Effects of *lft2* CRISPR-Cas9 injection on heart looping in F0 surface fish at 2 dpf. Q. Examples of mutations (underlined in green) in 4 CRISPR-Cas9 injected surface fish with mild axial phenotypes compared to a wild type control. Red arrows: sgRNA target. R. Bar graphs showing heart looping in genotyped CRISPR-CA9 injected surface fish with (top) or without (middle) mild axial phenotypes compared to controls from the same clutch (bottom).

The following experiments were performed to understand the role of *lft2* expression and its regulation in cavefish development. First, we asked if *lft2* downregulation is restricted to the LPM or is more general in cavefish embryos. Because *lft2* is also expressed during gastrulation [20, 21], *lft2*, and as a control *lft1*, expression was compared in surface fish and cavefish at 50% epiboly (Fig. 3I-L; N-P). *In situ* hybridization showed that *lft1* is expressed in the shield region of both surface fish and cavefish gastrulae (Fig. 3I-L), as described previously [22], and *lft2* is expressed in the germ ring in surface fish embryos (Fig. 3N) and cavefish embryos, although staining levels varied in individual cavefish gastrulae (Fig. 3O, P). The *in situ* hybridization results were confirmed by qPCR, which showed no significant differences in *lft2* or *lft1* mRNA levels between surface fish and cavefish gastrulae, although higher levels of *ndr1* mRNA were detected in cavefish (Fig. 3M), as demonstrated previously [23]. These results suggest that strong *lft2* downregulation is restricted to the cardiac field in the cavefish LPM. Second, to explore the mechanism of *lft2* downregulation, we PCR amplified and sequenced the four *lft2* exons in genomic DNA from individual male and female F61 PA-CF whose offspring showed high levels of reversed heart asymmetry (Fig. S4). No differences were uncovered in the nucleotide sequences of the *lft2* exons in cavefish of either sex, indicating that the *lft2* coding region is intact. The decrease in *lft2* expression during segmentation, but not during gastrulation, and the absence of coding mutations suggests that *lft2* downregulation is controlled by *cis*- or *trans*-acting changes in regulatory regions or by epigenetic mechanisms [24-26] in cavefish. Lastly, to investigate the role of *lft2* in heart laterality, surface fish with downregulated *lft2* expression were produced by CRISPR-Cas9 editing. In these experiments, surface fish eggs were injected with CRISPR-Cas9 and 2 sgRNAs, and larvae with minor axial defects were selected for genotyping (Fig 3Q). The remaining larvae were separated into groups with and without axial effects and were assayed for heart looping. We found increased proportions of larvae with altered heart looping in both groups of CRISPR-Cas9 larvae relative to controls from the same clutch of eggs (Fig. 3R). Although the shift to L-looped and un-looped hearts was modest, perhaps due to mosaic effects in the F0 CRISPR-edited embryos, the results support the possibility that *lft2* mutations can affect L-R axis determination. Genetic and molecular ablation of *lft2* also has minimal effects on L-R symmetry in zebrafish embryos [21, 27], suggesting that Nodal signaling in the absence of Lefty is functional but fragile. Fragility of Nodal signaling may also contribute to changing L-R symmetry in cavefish embryos with reduced *lft2* expression, perhaps through unknown genes involved in this process.

### Maternal Genetic Effects on Heart Looping Asymmetry

Based on the results of reciprocal hybridization experiments, the evolution of some cavefish traits is thought to be under maternal genetic control [22, 28]. Therefore, we asked whether cavefish heart looping changes are controlled by maternal or zygotic events. To distinguish between these possibilities, reciprocal hybridization experiments were performed in which a F61 PA-CF female was crossed with a male surface fish and a surface fish female was crossed with a F61 PA-CF male, and heart looping in the hybrid progeny was determined (Fig. 4A). As controls, the same surface fish and cavefish individuals used for reciprocal hybridization were crossed with each other. The results showed that heart laterality in the hybrids was dependent on the origin of the eggs: cavefish (female) X surface fish (male) hybrids produced heart laterality defects very similar to cavefish X cavefish controls (e. g. a mixture of D-looped, non-looped, and L-looped hearts), and surface fish (female) X cavefish (male) hybrids showed the same prevalence of D-looped hearts as the surface fish X surface fish progeny (Fig 4B). The results suggest that maternal genetic effects may be involved in evolutionary changes in cavefish heart laterality.

**Figure 4.**
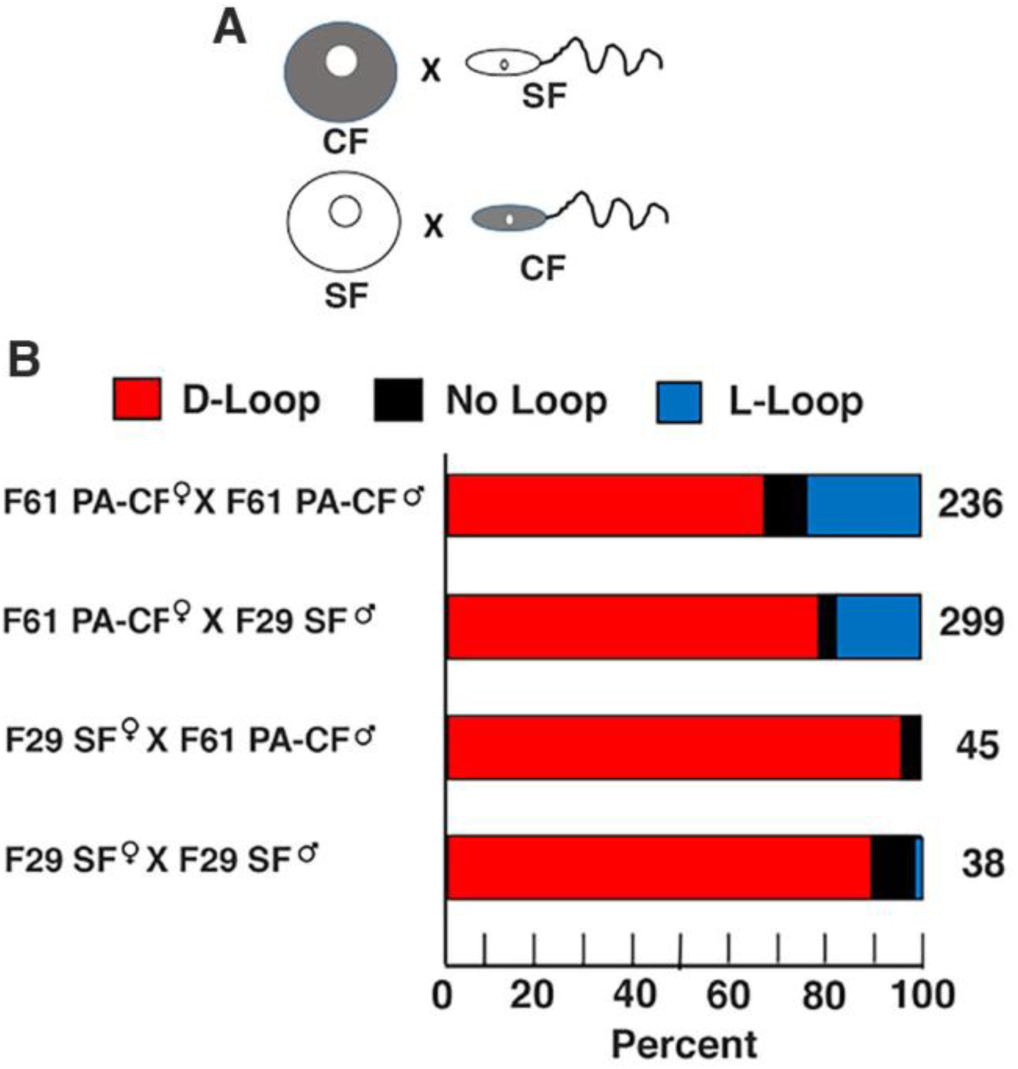
Maternal genetic effects on heart looping laterality. A. Reciprocal crosses between surface fish and cavefish gametes in both directions were used to assay for maternal genetic effects. B. Bar graphs showing the percentage of heart looping types in the F1 progeny of F61 cavefish x cavefish (top row), F61 cavefish female x F29 male (second from top row), F29 surface female x F61 cavefish male (second from bottom row), and F29 surface fish x F29 surface fish male (bottom row) crosses shown from top to bottom. Number of larvae analyzed are indicated at the right of each bar in B.

In conclusion, we have found that cavefish show natural changes in L-R axis determination, featuring reversals in the direction of the Nodal-Pitx2 signaling cascade, heart looping, and liver and pancreas polarity. These results suggest that the conserved L-R asymmetry of ancestral surface fish has been modified during cavefish evolution, and is to our knowledge the first instance of evolutionary changes of L-R patterning in a vertebrate. The L-R changes have no effects on survival, explaining their ability to be inherited in cavefish populations. The proportion of reversed organ asymmetry was greater in PA-CF than in TI-CF, consistent with more advanced cave related traits, including constructive traits and regressive eye and pigmentation traits, in the PA-CF population [7, 29, 30]. This suggests that L-R reversal may be related to cave adaptation, although the potential benefits, if any, and the evolutionary forces involved in this process, remain to be determined. Classic genetic studies demonstrated a maternal basis for L-R asymmetry of shell coiling in snails [31], and recent studies suggest that the maternally expressed *diaphanous* mRNA may be involved in this process [32, 33]. The possibility of a maternal origin of L-R symmetry has been more controversial in vertebrates [34], although a mutated maternal effect locus has been identified in zebrafish [35]. The reciprocal hybridization experiments described here support a maternal origin of L-R modifications in cavefish and suggest that *A. mexicanus* will provide an excellent model for determining the molecular basis of the first L-R symmetry breaking events during vertebrate development.

## Materials and Methods

### Animal Husbandry and Biological Procedures

*A. mexicanus* surface fish were originally collected at Nacimiento del Rio Choy, San Luis Potosi, Mexico, and Pachón (PA-CF) and Tinaja (TI-CF) cavefish were originally collected at La Cueva de El Pachón, Tamaulipas, Mexico and El Sótano de la Tinaja, San Luis Potosi, Mexico respectively. Families consisting of 10-20 individual male and female siblings of third generation progeny of wild captured fishes were raised in 40 L tanks at 22-23°C in a constant water flow system, fed a diet of TetraMin Pro flakes (Tetra Holding Inc, Blacksburg VA) and black worms (Eastern Aquatics, Lancaster, PA), and induced to spawn by excess feeding and gradual increase of water temperature to 25-26°C, as described previously [36]. In some cases, individual males and females were separated from a family, raised together, and subjected to paired mating. Reciprocal hybridization of cavefish females X surface fish males and surface fish females X cavefish males was carried out by *in vitro* fertilization as described previously [32]. Embryos and larvae were cultured at 23°C and fed living brine shrimp beginning at about 6-days post-fertilization (dpf). Animals were maintained and handled according to animal care guidelines of the University of Maryland, College Park (IACUC #R-NOV-18-59) (Project 1241065-1).

### Determination of Cardiac Laterality

The direction of cardiac tube jogging was determined visually by viewing larvae by viewing transparent larvae euthanized with 2 µg MS222 (Tricaine; Western Chemical Inc, Ferndale, CA) at 1.5-2 dpf. The direction of heart looping was determined at 3 dpf by staining with the myosin-heavy chain antibody MF-20 [11, 12] (Developmental Studies Hybridoma Bank, University Iowa, Iowa City, IA). For MF-20 staining, larvae were fixed with 4% paraformaldehyde (PFA) in PBS at 4°C overnight, washed three times with PBST (PBS, 0.5% Triton X-100), dehydrated by treatment with an increasing methanol series (25%, 50%, 75%) to 100% methanol, and stored at -20°C. Prior to antibody staining, the specimens were re-hydrated through a decreasing (75%, 50%, 25%) methanol series to PBST. The specimens were washed in chilled acetone, incubated in acetone for 7 min at -20°C, quickly rinsed in double distilled water, washed twice in PBST for 5 min, washed once with PBDT (PSA, 1% BSA, 1% DMSO, 0.5% Triton X-100), and blocked with 5% goat serum (Vector Laboratories, Burlingame, CA) in PBDT. Primary antibody staining (1:10 dilution) was done at 4°C overnight, and was followed by three washes with PBDT for 10 min and incubation with goat anti-mouse secondary antibody (1:500) (ThermoFisher, Waltham, MA) at 4°C overnight. The specimens stained with secondary antibody were washed in PBDT three times for 10 min, cleared in 50% and 75% glycerol, mounted in 75% glycerol, imaged, and photographed.

### Determination of Liver and Pancreas Laterality

The positioning of the liver and pancreas with respect to the middle line was determined in 60 hpf larvae by *in situ* hybridization (see below) for the *cystathionine ß-synthase a* (*cbsa*) gene. Previous studies have shown that *cbsa* is strongly expressed in the developing liver and pancreas at this stage of surface fish and cavefish development [13].

### Survival Analysis

Survival analysis was conducted on surface fish and PA-CF larvae separated into groups with D-looped, non-looped, or L-looped hearts. Cardiac looping was determined by visual inspection of 3 dpf anesthetized larvae viewed from the ventral side under a stereomicroscope. Larvae with different categories of heart looping were isolated in glass bowls in 50 ml of fish system water, fed brine shrimp beginning at 6 dpf, and counted daily. Counts were done on anesthetized larvae viewed under a stereomicroscope. Following counting, the larvae were rinsed several times in fish system water and returned to the bowls. Dead larvae were removed from the bowls daily. Significance was determined by the Cox proportional hazards model in R [37].

### Movie

PA-CF larvae at 3 dpf were mounted in 1.5% low melting point agarose. Movies were acquired using a Leica M205 stereo microscope with a Lecia DFC 7000 camera.

### *In situ* hybridization

The processing of specimens and procedures for *in situ* hybridization were carried out as described by Ma et al. [38]. Embryos were dechorionated manually using forceps, fixed in 4% PFA overnight, dehydrated in methanol, and stored at -20°C prior to *in situ* hybridization. The RNA probes used for *in situ hybridization* were prepared using the oligonucleotide primers (see Table S1) from *A. mexicanus* draft genome information [39]. After the completion of hybridization, the embryos were washed with PBST and incubated in BM Purple AP Substrate (Roche, Basel, Switzerland) at room temperature in the dark. After the signal developed, the reaction was terminated by rinsing the embryos in PBST. The embryos were processed through an increasing glycerol series in Phosphate Buffered Saline, imagining was done using an Olympus SZX12 microscope, and photography was carried out using a Zeiss Axioskop compound microscope

### RNA extraction and RT-PCR

Total RNA was extracted using TRI Reagent Solution (Life Technologies, Grand Island NY, USA), treated with RNase-free DNase (Qiagen, Frederick, MD) to remove traces of genomic DNA, and cDNA was synthesized using the SuperScriptTM III First-Strand Synthesis SuperMix Kit and random hexamer primers (ThermoFisher, Rockville, MD). Quantitative real time RT-PCR (qPCR) was done as described by Ma et al. [28] using the oligonucleotide primers shown in Table S2. To confirm the specificity of the primers, BLAST searches were done against the Ensembl Mexican tetra genome database. Dissociation curves were used to confirm the amplification of single PCR products. All reactions were replicated at least three times. For relative quantification, the ΔCt values for each gene were normalized and converted to 2^-ΔΔCt values, which represent the mean fold change of CF compared to SF mRNA levels. Statistical analysis using ΔCt values was conducted by Student’s t test.

### Sequencing of *lefty2* Genes

The *lft2* exon regions were amplified by PCR from genomic DNA isolated from tail fin clips of a surface fish and individual F61 PA-CF male and females using the Phusion High-Fidelity PCR Master Mix (New England Biolabs, Ipswich, MA) and the primers listed in Table S3. The PCR cycling conditions were 1 cycle of initial denaturation at 98°C for 30 sec, followed by 35 cycles each of denaturation (98°C for 8 sec), annealing (at 60°C for 15 sec), and extension (at 72°C for 15 sec) with a final elongation step at 72°C for 8 minutes. The PCR products were detected by gel electrophoresis and purified with the MinElute PCR Purification Kit (Qiagen, Valencia, CA, USA) and sequenced.

### CRISPR-Cas9 Gene Editing

To edit the *lft2* gene, 50pg of each sgRNA and 300pg Cas9 protein (Cas9 nuclease 2NLS, *S. pyrogenes*, Synthego Corp., Redwood City, CA) were co-injected into one-cell stage surface fish embryos. The two sgRNA were synthesized according to *lft2* sequence information (Ensembl ENSAMXT00000000477): the sequence of the first sgRNA was 5’-GCUUGCAGCACUGUCAAACA-3’, and the sequence of the second sgRNA was 5’-CAAACAGAAACAUGUGCAUG-3’. Microinjection was carried out as described previously [14, 38]. CRISPR-Cas9 injected and control un-injected embryos from the same clutch were cultured at 25 °C. At about 2 dpf, the injected larvae were viewed by microscopy, those with axial defects, such as twisted bodies or bent tails, were identified, and some of these were used to extract DNA and genotype the edited sites by nested PCR. For nested PCR, the flanking primers were 5’-GGCTCTAATGTGTCGTGCCT -3’ (forward) and 5’-ACACGATGACAAAACTACCCCT - 3’ (reverse), and the nested primers were -5’-TCTAGACGTGATGCAGGGGA -3’ (forward) and 5’-GCCTTAACATACCTATGCCAGC -3’ (reverse). The purified PCR products were assayed for genome targeting efficiency by sequencing. All of the remaining CRISPR-Cas9-sgRNA injected larvae and un-injected controls from the same clutch were assayed at 3 pdf for heart looping as described above.

## Acknowledgements

We thank Ruby Dessiatoun for animal care and Dr. Špela Gorički for assistance with statistics for survival curves. This research was supported by NIH grant EY024941 to W. R. J.

## Author Contributions

L.M., A.V.G., and W.R.J. conceived the project. L. M., M. N., J. S, and D. C. performed the research. W. R. J. and L. M. wrote the manuscript with input from other authors.

## References

1. Nakamura, T., Hamada, H. (2012). Left-right patterning: conserved and divergent mechanisms. Development 139, 3257–3262.

2. Blum, M., Feistel, K., Thumberger, T, and Schweickert, A. (2014). The evolution and conservation of left-right patterning mechanisms. Development 141, 1603–1613.

3. Grimes, D. T., and Burdine, R. D. (2017). Left-right patterning: breaking symmetry to asymmetric morphogenesis. Trends Genet. 33, 616–628.

4. Montague, T., Gagnon, J. A., and Schier, A. (2018). Conserved regulation of Nodal-mediated left-right patterning in zebrafish and mouse. Development 145, dev171090 doi: 10.1242/dev.171090

5. Chen, J-C., van Bebber, F., Goldstein, A. M., Serluca, F. C., Jackson, D., Childs, S., Serbedzija, G., Warren, K. S., Mably, J. D., Lindahl, P., Mayer, A., Haffter, P., and Fishman, M. C. (2001). Genetic steps to organ laterality in zebrafish. Comp. Func. Genom. 2, 60–68.

6. Shiraishi, I., and Ichikawa, H. (2012). Human heterotaxy syndrome: from molecular genetics to clinical features, management, and prognosis. Circ. J. 76, 2066–2075.

7. Jeffery, W. R. (2001). Cavefish as a model system in evolutionary developmental biology. Dev. Biol. 231, 1–12.

8. Herman, A., Brandvain, Y. Weagley, J., Jeffery, W. R., Keene, A. C., Kono, T. J. Y., Bilandžija, H., Borowsky, R., Espinasa, L. O’Quin, K., Ornelas-García, C. P., Yoshizawa, M., Carlson, M., Maldonado, E., Gross, J. B., Cartwright, R. A., Rohner, N., Warren, W. C., and McGaugh, S. E. (2018). The role of gene flow in rapid and repeated evolution of cave-related traits in Mexican tetra, Astyanax mexicanus. Mol. Ecol. 27, 4397–4416.

9. Powers, A. K., Davis, E. M., Kaplan, S. A., and Gross, J. B. (2017). Cranial asymmetry arises later in the life history of the blind cavefish, *Astyanax mexicanus*. PLoS ONE 12(5): e0177419. https://doi.org/10.1371/journal.pone.0177419

10. Powers, A. K., Boggs, T. E., and Gross, J. B. (2018). Cranial neuromast position pre-figures developmental patterning of the suborbital bone series in *Astyanax* cave-and surface-dwelling fish. Dev. Biol. 441, 252–261.

11. Wolpert, L, Beddington, R., Jessell, T., Lawrence, P., Meyerowitz, E, and Smith, J. (2002). Principles of Development. Oxford University Press, Oxford.

12. Bader, D., Masaki, T., and Fischman, D. A. (1982). Immunochemical analysis of myosin heavy chain during myogenesis in vitro and in vivo. J. Cell Biol. 95, 763–770.

13. Berdougo, E., Coleman, H., Lee, D. H., Stainier, D. Y., and Yelon, D. (2003). Mutation of weak atrium/atrial myosin heavy chain disrupts atrial function and influences ventricular morphogenesis in zebrafish. Development 130, 6121–6129.

14. Ma, L. Gore, A. V., Castranova, D., Shi, J., Ng, M., Tomins, K. A., van der Weele, C. M., Weinstein, B. M., and Jeffery, W. R. (2020). A hypomorphic *cystathionine ß-synthase* gene contributes to cavefish eye loss by disrupting optic vasculature. Nature Commun. (in press).

15. Field, H. A., Ober, E. A., Roeser, T., and Stanier, D. Y. R. (2003). Formation of the digestive system in zebrafish. I. liver morphogenesis. Dev. Biol. 253, 279–290.

16. Essner, J. J., Amack, J. D., Nyholm, M. K., Harris, E. B., and Yost, H. J. (2005). Kupffer’s vesicle is a ciliated organ of asymmetry in the zebrafish embryo that initiates left-right development of the brain, heart, and gut. Development 132, 1247–1260

17. Long, S., Ahmad, N., and Rebagliati, M. (2003). The zebrafish nodal-related gene southpaw is required for visceral and diencephalic left-right symmetry. Development 130, 2303–2316.

18. Meno, C., Shimono, A., Saitoh, Y., Yashiro, K., Mochida, K., Ohishi, S., Noji, S. Kondoh, and Hamamda, H. (1998). lefty-1 is required for left-right determination as a regulator of lefty-2 and nodal. Cell 94, 287–297.

19. Bisgrove, B., Essner, J., and Yost, H. J. (1999). Regulation of midline development by antagonism of lefty and nodal signaling. Development 126, 3253–3262.

20. Lenhart, K. F., Lin, S-Y., Titus, T. A., Postlethweit, J. N., and Burdine, R. D. (2011). Two additional midline barriers function with midline *lefty1* expression to maintain asymmetric Nodal signaling during left-right axis specification in zebrafish. Development 138, 4405–4410.

21. Van Boxtel, A. L., Econonmou, A. D., Heliot, C., and Hill, C. S. (2017). Long range signaling activation separate the mesoderm and endoderm lineages. Dev. Cell 44, 179–191.

22. Rogers, K. W., Lord, D. D., Gagnon, J. A., Pauli, A., Zimmerman, S., Aksel, D. C., Reyon, D., Tsai, S. Q., Joung, J. K., and Schier, A. F. (2017). Nodal patterning without Lefty inhibitory feedback is functional but fragile. eLife 6, e28785 DOI: 10.7554/eLife.28785

22. Torres-Paz, G., LeClercq, J., and Rétaux, S. (2019). Maternally regulated gastrulation as a source for variation contributing to cavefish forebrain evolution. eLife 8, e50160 DOI: 10.7554/eLife.50160

23. Ren, X., Hamilton, N., Muller, F., and Yamamoto, Y. (2018). Cellular rearrangement of the prechordal plate contributes to eye degeneration in the cavefish. Dev. Biol. 441, 221–234.

24. Gore, A. V., Tomins, K. A., Iben, J., Ma, L., Castranova, D., Davis, A., Parkhurst, A., Jeffery, W. R., and Weinstein, B. M. (2018). An epigenetic mechanism for cavefish eye degeneration. Nature Ecol. Evol. 2, 1155–1160. doi: 10.1038/s41559-018-0569-4

25. Wang, L., Liu, Z., Lin, H., Ma, D., Tao, D., and Liu, F. (2017). Epigenetic regulation of left-right asymmetry by DNA methylation. EMBO J. 36, 2987–2997.

26. Dai, H-Q., Wang, B-A., Yang, L., Chen, J-J., Zhu, G-C., Sun, M-L., Ge, H., Wang, R., Chapman, D. L., Tang, F., Sun, X., and Xu, G-L. (2016). TET-mediated DNA methylation controls gastrulation by regulating Lefty-Nodal signalling. Nature 538, 528–532.

27. Meno, C., Takeuchi, J., Sakuma, R., Koshiba-Takeuchi, K., Ohsishi, S., Saijoh, Y., Miyazaka, J., ten Dijke, P., Ogura, T., and Hamada, H. (2001). Diffusion of Nodal signaling activity in the absence of the feedback inhibitor Lefty2. Dev. Cell 1, 127–138.

28. Ma, L, Strickler, A. E., Parkhurst, A., Yoshizawa, Y., Shi, J., and Jeffery, W. R. (2018). Maternal genetic effects in *Astyanax* cavefish development. Dev. Biol. 441: 209-220.

29. Jeffery, W. R. (2009). Evolution and development in the cavefish *Astyanax*. Curr. Top. Dev. Biol. 86, 191–221.

30. Keene, A., Yoshizawa, M., and McGaugh, S. (2016). Biology and evolution of the Mexican cavefish. New York; Elsevier; 2016.

31. Sturtevant, A. H. (1923). Inheritance of the direction of coiling in *Limnaea*. Science 58, 269–270.

32. Davison, A., McDowell, G. S., Holden, J. M., Johnson, H. F., Koutsovoulos, G., Liu, M. M., Hulpiau D., Roy, F. J., Wade, C. M, Banerjee, R., Yang, F., Chiba, S., Davey, J. W., Jackson, D. J., Levin, M., and Blaxter, M. L. Formin is associated with left-right asymmetry in the pond snail and frog. Curr. Biol. 26, 654–660.

33. Abe, M., and Kuroda, R. (2019). The development of CRISPR for a mollusk establishes the formin *Lsdia1* as the long-sought gene for snail dexral/sinistral coiling. Development 146, dev175976 doi: 10.1242/dev.175976

34. Schweickert, A., Walentek, P., Thumberger, T., and Danilchek, M. (2012). Linking early determinants and cilia-driven leftward flow in left-right axis specification of *Xenopus laevis*: A theoretical approach. Differentiation 83, S67–S77

35. Domenichini, A., Dadda, M., Facchin, L., Bisazza, A, and Argenton, F. (2011). Isolation and genetic characterization of *Mother-of-Snow-White*, a maternal effect allele affecting laterality and lateralized behaviors in zebrafish. PLoS ONE 6, (10): e25972. https://doi.org/10.1371/journal.pone.0025972

36. Jeffery, W. R., Strickler, A. G., Guiney, S., Heyser, D., and Tomarev, S. I. (2000). *Prox 1* in eye degeneration and sensory compensation during development and evolution of the cavefish *Astyanax*. Dev. Genes. Evol. 210, 223–230.

37. Cox, D. R. (1972). Regression models and life tables (with discussion). J. R. Statist. Soc. B 34, 187–220.

38. Ma, L., Parkhurst, A., and Jeffery, W. R. (2014). The role of a lens survival pathway including *sox2* and *αA-crystallin* in the evolution of cavefish eye degeneration. EvoDevo 5, 28 doi:10.1186/2041-9139-5-28.

39. McGaugh, S. E., Gross, J. B., Aken, B., Blin, M., Borowsky, R., Chalopin, D., Hinaux, H., Jeffery, W. R., Keene, A., Ma, L., Minx, P., Murphy, D, O’Quin K. E., Rétaux, S., Rohner, N., Searle, S. M. J., Stahl, B., Tabin, C., Volff, J. N., Yoshizawa, M., and Warren, W. (2014). The cavefish genome reveals candidate genes for eye loss. Nature Commun. 5, doi.10.1038/natcommun630

