## Supplementary Files for "Evolutionary Changes in Left-Right Visceral Asymmetry in *Astyanax* Cavefish"

**Supplementary Movie.**

Video showing beating of right (D) lopped and left (L) looped hearts in Pachón cavefish at 3 days post-fertilization.

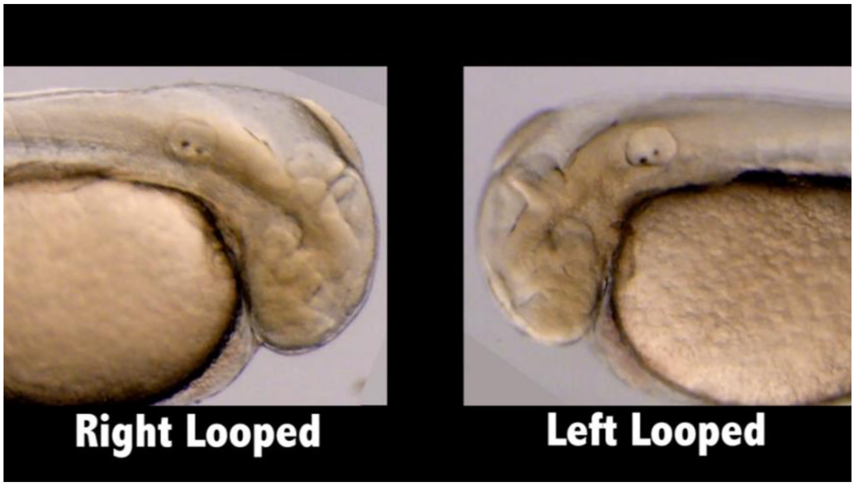


**Supplementary Figures and Legends**.

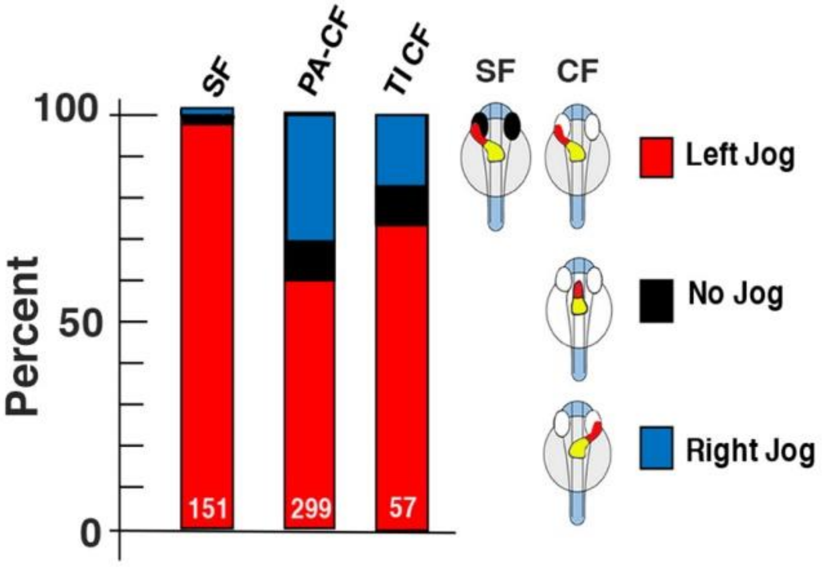


Figure S1. Bar graph showing the proportion of left jogged, non-jogged, and right jogged cardiac tubes in surface fish (SF), Pachón cavefish (PA-CF), and Tinaja cavefish (TI-CF) at 1.5 dpf. N is shown at the bottom of each bar.

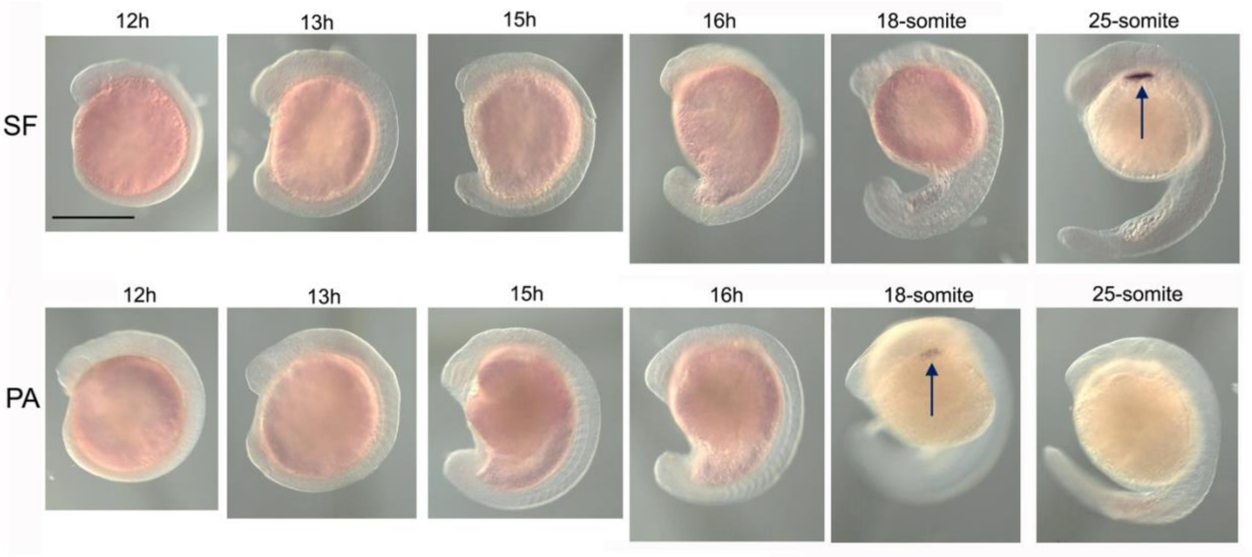


Figure S2. Determination of *lft2* gene expression in left lateral plate mesoderm (LM) of surface fish and cavefish (F 61 PA-CF) by *in situ* hybridization between the 12 hour (h) and 25-somite stages. The *lft2* gene is expressed strongly in the anterior left LPM at the

25-somite stage in surface fish embryos (arrow) but weakly in cavefish embryos at the 18-somite stage (arrow).  Scale bar is 200 µm; magnification is the same in all frames.

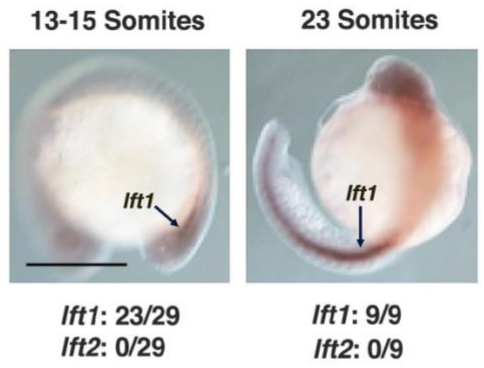


Figure S3. Double *in situ* hybridization with *lft1* and *lft2* probes. Stages shown at top of each frame.  Number of *lft1* or *lft2* stained individuals per total number shown on bottom of frames. Scale bar is 200 µm; magnification is the same in all frames.

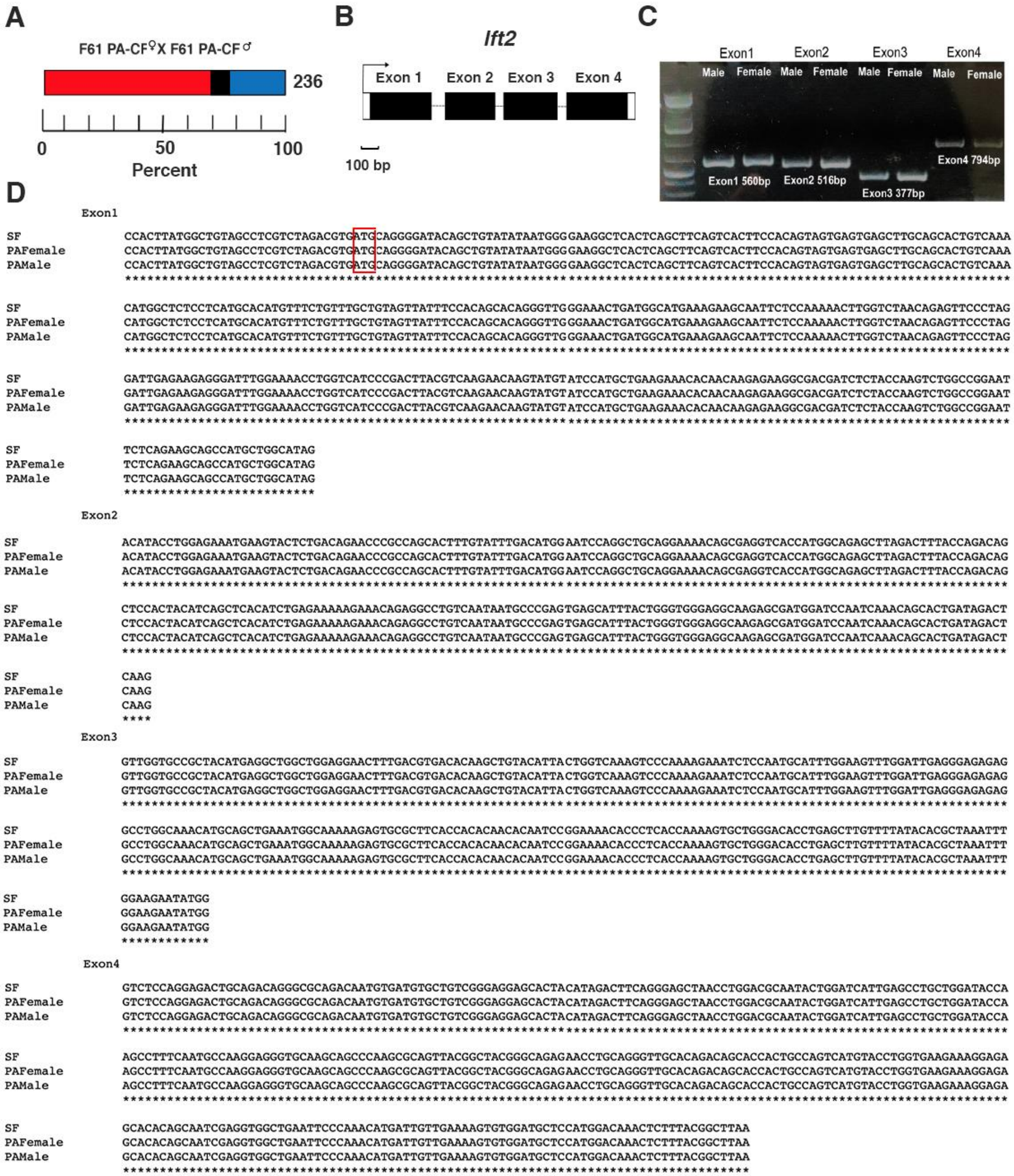


Figure S4. Survey of cavefish *lft2* genomic DNA for coding region changes. A. Comparison of numbers of embryos with D-looping hearts, no looping hearts, and left looping hearts in the progeny of a single surface fish x surface fish and a cavefish x cavefish (both F61 PA-CF) cross. B. Exon-intron organization of the *Astyanax lft2* gene. C. PCR amplification of 4 *lft2* exons from male and female F61 PA-CF individuals. Exons shown as red outlined bars; bars showing 5' UTR in exon 1 is not filled. Introns shown as lines. D. PCR amplification of *lft2* exons from male and female individuals used in the cavefish x cavefish cross in A. D. Alignment of *lft2* exon sequences obtained from the male and female individuals used in the cavefish x cavefish cross in A with the surface fish *lft2* exon sequence. Red box: ATG translation start site. Asterisks: identical nucleotides.

### Supplementary Tables

Supplementary Table 1. Oligonucleotide primers used to amplify gene sequences for preparation of RNA probes for *in situ* hybridization.

| Gene | Primers |
|---|---|
| *spaw* | Forward: TTTAACGTGACCGCTCTGCT<br>Reverse: TGCATGTAGGCGTGATTGGT |
| *lefty1* | Forward: CAGGACCCCAGCGATAACTC<br>Reverse: GCCGCACTTCTCCACTATCA |
| *lefty2* | Forward: GGCAAAAAGAGTGCGCTTCA<br>Reverse: TGTCCATGGAGCATCCACAC |
| *pitx2* | Forward: CCCAAAATGGACGCAAAGGG<br>Reverse: TATGGTGGCAATTGCAGGGT |
| *cbsa* | Forward: CGCATGCTCATCAGAGACGA<br>Reverse: GGCAAAGTGATCCGTCTCCA |

Supplementary Table 2. Oligonucleotide primers used in RT-PCR.

| Gene name | Froward primers | Reverse primers |
|---|---|---|
| *spaw* | CGCTAAAGACTGTCATCAGGTTG | AACAACAGCCCGTTTGGTTG |
| *pitx2* | CTACACACCCCCTTAGCCAT | GTCTTTATCTGCGCACTCGG |
| *lft1* | GACCCCAGCGATAACTCACT | CTGCAGCACTGACCCTGA |
| *lft2* | GAATCAGTCTTCGCGTTATTTCC | GACGTAAGTCGGGATGACCA |
| *ndr1* | ACCCTAAGCGATACAATGCCT | AGCTTCAGAAGACTCTGCATGT |
| *ndr2* | CACGCCTACATGCAGAGTC | TCTCGCCGTTCTCGTAGTAG |
| *gapdh* | TCCTGAACTCAATGGCAAGC | TTCTCCAAGCGGACAGTCAA |

Supplementary Table 3. Oligonucleotide primers used to amplify *lft2* exons by PCR.

| *lft2* Exon | Forward primers | Reverse primers |
|---|---|---|
| 1 | GCACAGTTTGGGCAACAGAG | AAAGCAGAGCCTTAACATACCT |
| 2 | GCATAGGTATGTTAAGGCTCTGC | CACGATGACAAAACTACCCCT |
| 3 | TCAGGGGTAGTTTTGTCATCGT | ACACACACCTCAACATTACCTCA |
| 4 | TGAGGTAATGTTGAGGTGTGTGT | AACTGTCGAGTGTTGCCGTA |
